# Twenty novel nsSNPs may affect *FLT3* gene leading to Acute Myeloid Leukemia (AML) using in silico analysis

**DOI:** 10.1101/2023.06.24.546344

**Authors:** Tarig Alsheikh, Tebyan Ameer, Ahmed NjmEldin, Dalia Omer, Abubaker Aghbash, Hadil Suliman, Zeinab Abdalmonem, Howiada Hamad, Saif Eldowla A. ayoub, Mohammed A. Hassan

## Abstract

**Background**: Mutations within the FMS-like tyrosine kinase 3 (*FLT3*) gene represent one of the most common genetic alteration that disturb intracellular signaling networks with a key role in leukemia pathogenesis. laboratory studies considerable obstacle to identify functional SNPs in a specific gene. Thus, the “in silico” technique is possible now to carry out research investigations without the need for extensive lab work. **Methodology**: data retrieved from NCBI database and different algorithm used to analyse nsSNPs which they are: SIFT, Polyphen-2, Provean, SNAP2, P-Mut, I-Mutant, Project Hope, Raptor X, PolymiRTS and Gene MANIA. **Result:** Our study reveals twenty novel SNPs regarded to be the most damaging SNPs that affect structure and function of *FLT3* gene using different bioinformatics algorithm. **Conclusion**: This study revealed 20 damaging SNPs considered to be novel nsSNP in *FLT3* gene that leads to AML, by using different algorithms. Additionally, 69 functional classes were predicted in 12 SNPs in the 3’UTR, among them, 31 alleles disrupted a conserved miRNA site and 37 derived alleles created a new site of miRNA. This might result in the de regulation of the gene function. These results could be valuable for molecular studying, diagnosis and treatment of AML patients.

## INTRODUCTION

Acute myeloid leukemia (AML) is a heterogeneous hematologic malignancy characterized by infiltration of the bone marrow, blood, and other tissues by proliferative, clonal, abnormally differentiated, and occasionally poorly differentiated cells of the hematopoietic system (1). It is the most frequent form of acute leukemia in adults, and it causes the most leukemia-related deaths in the United States each year (2).

AML is classified according to the World Health Organization (WHO) Classification of Tumours of Haematopoietic and Lymphoid Tissues. The major categories of the classification include AML with recurrent genetic abnormalities, AML with myelodysplasia-related changes, therapy-related AML, and AML not otherwise specified (1).

AML is the result of distinct but cooperating genetic mutations that promote proliferation, survival and impair differentiation and apoptosis including mutations of *FLT3*, ALM, oncogenic RAS and PTPN11, and the BCR/ABL (3).

Mutations within the FMS-like tyrosine kinase3 (*FLT3*) gene represent one of the most common genetic alteration that disturb intracellular signaling networks with a key role in leukemia pathogenesis (4)

*FLT3* gene encodes protein called FMS-like tyrosine kinase3 (*FLT3*), which is part of a family of proteins called receptor tyrosine kinases type III (RTKs) that is originally expressed on hematopoietic progenitor cells (5) and it is play an important role in normal hematopoietic stem cells function (6).

The *FLT3* protein is found in the outer membrane of certain cell types where is a specific protein called *FLT3* ligand, can bind to it, Upon this binding, *FLT3* receptors are activated and dimerize, leading to activation of a series of proteins inside the cell, this multiple signaling pathways control cellular growth, proliferation, and cell survival (7).

The *FLT3* gene is located on chromosome 13 (13q12), which is consists of 24 exons and covers approximately 96 kb and encodes a 993–amino acid protein in human (8).

There are two major types of *FLT3* mutations internal tandem duplication mutation in the juxta-membrane domain (*FLT3*-ITD) and point mutation or deletion in the tyrosine kinase domain (*FLT3*-TKD) (9). Both types of mutations constitutively activate *FLT3* tyrosine kinase activity, that gain-of-function mutations of *FLT3* associates with leukemogenesis and poor prognosis in AML patients (4, 10)

The influence of mutations in these gene on AML patients remain controversial, and the clinical impact of these mutations remains unclear (11).

Identification of the functional SNPs in a particular gene using laboratory investigations still presents a significant challenge. As a result, research inquiries can now be conducted using the "in silico" technique without the requirement for intensive lab work. Too yet, computational study has not revealed the single nucleotide polymorphism for the *FLT3* gene. Therefore, the aim of this study is to perform a computational analysis of the SNPs in the *FLT3* gene, different algorithms like SIFT, PROVEAN, PolyPhen 2 and SNAP2 were used to identify the most damaging SNPs and the effect they may impose on protein structure and function.

## MATERIALS AND METHODS

Data on *FLT3* gene was obtained from the national center for biological information (NCBI) website (https://www.ncbi.nlm.nih.gov/) and the SNPs (single nucleotide polymorphisms) information was retrieved from NCBI SNPs database dbSNP (https://www.ncbi.nlm.nih.gov/snp/). The gene ID and sequence was obtained from Uiprot (https://www.uniprot.org/). Analysis of the SNPs was done according to figure 1.

**Figure (1):**
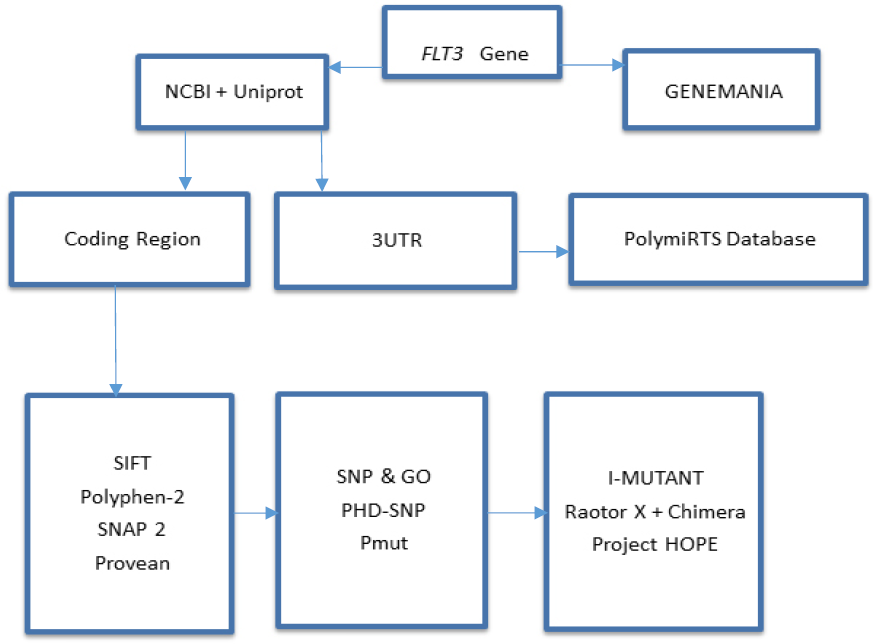
Diagram demonstrates *FLT3* gene work flow using different bioinformatics tool.

### SIFT

)Sorting Intolerant from Tolerant) is a sequence homology-based tool that distinguishes intolerant from tolerant amino acid substitutions and predicts whether an amino acid substitution in a protein will result in phenotypic change, taking into account the position of the mutation and the type of amino acid change. SIFT selects similar proteins and obtains an alignment of these proteins with the query when a protein sequence is submitted. SIFT calculates the chance that an amino acid at a location is tolerated based on the type of amino acids which is change in the appearing at each place in the alignment, conditional on the most frequent amino acid being tolerated. The replacement is projected to be detrimental if this normalized value is smaller than a cut-off. The method predicts intolerant or harmful amino acid substitutions for SIFT values less than 0.05, whereas scores more than 0.05 are considered tolerant. (12). (http://sift.bii.a-star.edu.sg/)

### PolyPhen-2

Polymorphism Phenotyping v.2 (http://genetics.bwh.harvard.edu/pph2/) is a tool that uses simple physical and comparative considerations to forecast the likely influence of an amino acid change on the structure and function of a human protein. The sequence submission enables searching for a single amino acid alteration or a coding, nonsynonymous SNP that has been annotated in the SNP database. It computes the difference between the PSIC scores of the two variants and creates position-specific independent count (PSIC) scores for each of the two variations. The greater the difference in PSIC scores, the greater the functional effect of a certain amino acid alteration is predicted to be. PolyPhen scores were divided into three categories: likely harmful (0.95–1), maybe harmful (0.7–0.95), and benign (0.00–0.31) (13).

### Provean

The Protein Variation Effect Analyzer (http://provean.jcvi.org/index.php) software predicts whether an amino acid substitution affects the protein’s biological activity. Provean is helpful for filtering sequence variants to find non-synonymous variants that are likely to have functional significance. (14).

### SNAP2

SNAP2 is a learned classifier that is based on a machine learning device called "neural network" and is available at (https://rostlab.org/owiki/index.php/Snap2). It considers a range of sequence and variant properties to distinguish between effect and neutral variants/non-synonymous SNPs. The evolutionary information extracted from an artificially generated multiple sequence alignment is the most crucial input signal for the prediction. Structures such as anticipated secondary structure and solvent accessibility are also taken into account. Annotation (i.e. known functional residues, pattern, regions) of the sequence or close homologs is pulled in if it is available. SNAP2 has an 82 percent two-state accuracy (effect/neutral) (15).

### PHD-SNP

(http://snps.biofold.org/phd-snp/phd-snp.html) predicts deleterious single nucleotide polymorphisms in humans. The working premise is a Support Vector Machines (SVMs)-based technique that uses sequence information to predict disease-associated nsSNPs. The associated mutation is classified as either disease-related (Disease) or neutral polymorphism (Polymorphism) (Neutral) (16).

### SNP and GO

It is a service that predicts single point protein mutations that are likely to be important in the development of human diseases (16). (https://snps-and-go.biocomp.unibo.it/snps-and-go/)

### Pmut

Available at (http://mmb.irbbarcelona.org/PMut/),Pmut is based on the use of many types of sequence information to flag mutations, as well as neural networks to analyse the data. It produces a straightforward result: a yes/no response, as well as a dependability index (17).

### IMUTANT

Imutant is a set of support vector machine-based predictors that have been integrated into a one-of-a-kind web server. It allows users to anticipate protein stability changes based on single site mutations using only the protein sequence or, if available, the protein structure. It also allows for the prediction of human harmful SNPs solely based on the protein sequence. It is available at (http://gpcr.biocomp.unibo.it/∼emidio/I-Mutant3.0/old/IntroIMutant3.0_help.html) (18).

### Project Hope

Online software is available at: (http://www.cmbi.ru.nl/hope/method/). It’s an online application that allows users to input sequences and mutations. The software gathers structural data from a variety of sources, including 3D protein structure computations, UniProt sequence annotations, and other software prediction. It integrates this data to produce a report on the impact of a specific mutation on the protein structure. HOPE will demonstrate the impact of that mutation in such a way that even persons with no prior knowledge of bioinformatics will be able to comprehend it. User can enter a protein sequence (FASTA or not) or an accession code for the protein of interest with a simple mouse click, the user can designate the altered residue in the next step. User can simply click on one of the other 19 amino acid types that will become the mutant residue in the final phase, and a comprehensive report will be provided (19).

### UCSF Chimera (University of California at San Francisco)

UCSF Chimera (https://www.cgl.ucsf.edu/chimera/) is a flexible tool for visualizing and analyzing molecular structures and related data, including a density maps, supramolecular assemblies, sequence alignments, docking findings, trajectories, and conformational ensembles. It is possible to create high-quality photos and animations. Chimera comes with comprehensive documentation and a number of tutorials. Chimera is being developed by the National Institutes of Health-funded Resource for Biocomputing, Visualization, and Informatics (RBVI). (P41-GM103311) (20).

### PolymiRTS

PolymiRTS is a software used to predict 3UTR (un-translated region) polymorphism in microRNAs and their target sites available at (http://compbio.uthsc.edu/miRSNP/). It’s a database of naturally occurring DNA variants in the seed region and target regions of mocriRNAs (miRNAs). MicroRNAs bind to the transcripts of protein-coding genes, causing mRNA instability or translational suppression. The PolymiRTS database was produced by screening 3UTRs of mRNAs in human and mouse for SNPs in miRNA target sites, which may impact miRNA-mRNA interaction and thereby miRNA-mediated gene regulation. The influence of polymorphism on gene expression and phenotypes is then determined, and the database is linked. Polymorphism in target sites that has been supported by a number of experimental approaches, as well as polymorphism in miRNA seed regions, are all included in the PolymiRTS data source (21).

### GeneMANIA

It is gene interaction software that uses a vast number of functional association data to locate other genes that are associated to a set of input genes. Protein and genetic relationships, pathways, co-expression, co-localization and protein domain similarity are all examples of association data. GeneMANIA can also be used to discover new members of a pathway or complex, locate extra genes that your screen may have missed or discover novel genes that perform a specific function, such as protein kinases. available at (https://genemania.org/) (22).

## RESULT

### Data Mining

Data of human *FLT3* gene were collected from the National Center for Biological Information (NCBI) website (23). The SNP information (SNP ID) of the *FLT3* gene was retrieved from the NCBI dbSNP (http://www.ncbi.nlm.nih.gov/snp/) and the protein ID and its sequence was collected from Swiss-Prot databases with the accession number: (P36888). (http://expasy.org/) (24).

### Functional analysis of nsSNP effect on *FLT3* using SIFT, Polyphen2, PROVEANand SNAP2 softwares

**Table (1):** Damaging (Deleterious) missense SNPs on *FLT3* associated variations predicted by SIFT, PolyPhen2, Provean, and SNAP2 softwares:

| dbSNP rs# | SNPs | SIFT prediction | SIFT score | Polyphen2 prediction | Polyphen2 score | PROVEAN score | PROVEAN prediction | SNAP2 prediction | SNAP2 score |
| --- | --- | --- | --- | --- | --- | --- | --- | --- | --- |
| rs764288814 | C231Y | D | 0 | probably damaging | 1 | -4.61 | Deleterious | effect | 79 |
|  | F349C | D | 0 | probably damaging | 1 | -6.1 | Deleterious | effect | 43 |
| rs1041412646 | W383C | D | 0 | probably damaging | 1 | -7.99 | Deleterious | effect | 78 |
| rs1442266284 | G619D | D | 0 | probably damaging | 1 | -6.54 | Deleterious | effect | 87 |
| rs780941129 | V624A | D | 0 | probably damaging | 1 | -3.73 | Deleterious | effect | 50 |
| rs1276477642 | V643D | D | 0 | probably damaging | 1 | -6.54 | Deleterious | effect | 75 |
| rs887807138 | C681Y | D | 0 | probably damaging | 1 | -10.6 | Deleterious | effect | 76 |
| rs887807138 | C681S | D | 0 | probably damaging | 1 | -9.61 | Deleterious | effect | 68 |
| rs1221467960 | C694R | D | 0 | probably damaging | 1 | -11.5 | Deleterious | effect | 81 |
| rs774511256 | Y702C | D | 0 | probably damaging | 1 | -6.61 | Deleterious | effect | 51 |
| rs1474119276 | D787Y | D | 0 | probably damaging | 1 | -8.42 | Deleterious | effect | 54 |
| rs568745490 | E858K | D | 0 | probably damaging | 1 | -3.67 | Deleterious | effect | 89 |
| rs774360726 | L860P | D | 0 | probably damaging | 1 | -5.03 | Deleterious | effect | 51 |
| rs759638195 | Y865C | D | 0 | probably<br>damaging | 1 | -8.24 | Deleterious | effect | 61 |
| rs868802040 | Y874C | D | 0 | probably<br>damaging | 1 | -7.83 | Deleterious | effect | 11 |
| rs776952835 | E880A | D | 0 | probably<br>damaging | 1 | -5.49 | Deleterious | effect | 82 |
| rs1423011599 | G885V | D | 0 | probably<br>damaging | 1 | -8.26 | Deleterious | effect | 82 |
| rs142796469 | P890R | D | 0 | probably<br>damaging | 1 | -8.23 | Deleterious | effect | 10 |
| rs200909894 | M908V | D | 0 | probably<br>damaging | 1 | -3.49 | Deleterious | effect | 25 |
| rs974980406 | M922T | D | 0 | probably<br>damaging | 1 | -5.28 | Deleterious | effect | 69 |
| rs1361689773 | C925Y | D | 0 | probably<br>damaging | 1 | -9.77 | Deleterious | effect | 68 |

### Disease effect & stability analysis of nsSNP on*DNMT3A* using P-Mut & I-Mutant softwares respectively

**Table (2):** Disease effect & stability analysis of nsSNPs associated variations predicted by P-Mut & I-Mutant version 3.0 respectively (*RI: Reliability Index):

| dbSNP rs# | Amino<br>change | SNP&GO<br>Prediction | RI | SNP&GO<br>Score | PHD-SNP<br>Prediction | RI | PHD-SNP<br>Score | I-MUTANT<br>Prediction | I-MUTANT<br>Score |
| --- | --- | --- | --- | --- | --- | --- | --- | --- | --- |
| rs764288814 | C231Y | Disease | 7 | 0.84 | Disease | 9 | 0.94 | Decrease | -0.36 |
| rs537215681 | F349C | Disease | 5 | 0.734 | Disease | 7 | 0.827 | Decrease | -1.73 |
| rs1041412646 | W383C | Disease | 3 | 0.659 | Disease | 8 | 0.876 | Decrease | -1.51 |
| rs1442266284 | G619D | Disease | 6 | 0.823 | Disease | 9 | 0.927 | Decrease | -0.92 |
| rs780941129 | V624A | Disease | 1 | 0.529 | Disease | 5 | 0.728 | Decrease | -1.81 |
| rs1276477642 | V643D | Disease | 5 | 0.775 | Disease | 8 | 0.886 | Decrease | -1.2 |
| rs887807138 | C681Y | Disease | 7 | 0.851 | Disease | 9 | 0.949 | Decrease | -0.42 |
| rs887807138 | C681S | Disease | 3 | 0.665 | Disease | 7 | 0.848 | Decrease | -0.96 |
| rs1221467960 | C694R | Disease | 6 | 0.793 | Disease | 8 | 0.887 | Decrease | -0.37 |
| rs774511256 | Y702C | Disease | 4 | 0.695 | Disease | 7 | 0.853 | Decrease | -1.32 |
| rs1474119276 | D787Y | Disease | 5 | 0.766 | Disease | 9 | 0.931 | Decrease | 0 |
| rs568745490 | E858K | Disease | 6 | 0.823 | Disease | 8 | 0.876 | Decrease | -0.6 |
| rs774360726 | L860P | Disease | 6 | 0.777 | Disease | 8 | 0.889 | Decrease | -1.6 |
| rs759638195 | Y865C | Disease | 5 | 0.752 | Disease | 8 | 0.887 | Decrease | -1.11 |
| rs868802040 | Y874C | Disease | 5 | 0.772 | Disease | 8 | 0.913 | Decrease | -1.11 |
| rs776952835 | E880A | Disease | 1 | 0.565 | Disease | 7 | 0.834 | Decrease | -0.82 |
| rs1423011599 | G885V | Disease | 7 | 0.855 | Disease | 9 | 0.935 | Decrease | -0.45 |
| rs142796469 | P890R | Disease | 5 | 0.763 | Disease | 8 | 0.91 | Decrease | -1.32 |
| rs200909894 | M908V | Disease | 1 | 0.564 | Disease | 7 | 0.848 | Decrease | -0.85 |
| rs974980406 | M922T | Disease | 5 | 0.769 | Disease | 8 | 0.881 | Decrease | -1 |
| rs1361689773 | C925Y | Disease | 7 | 0.848 | Disease | 9 | 0.947 | Increase | -0.25 |

### Modeling of amino acid substitution effects on protein structure using Project Hope and Chimera Softwares

**Figure (2):**
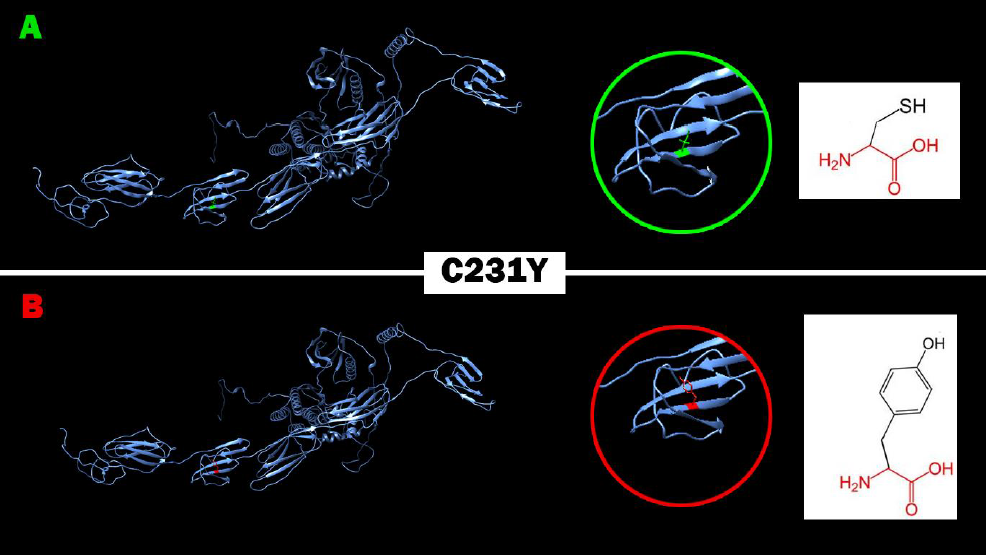
Effect of C231Y (rs764288814) SNP on protein structure in which Cysteine (Green residue) mutated into Tyrosine (Red residue) at position 231.

**Figure (3):**
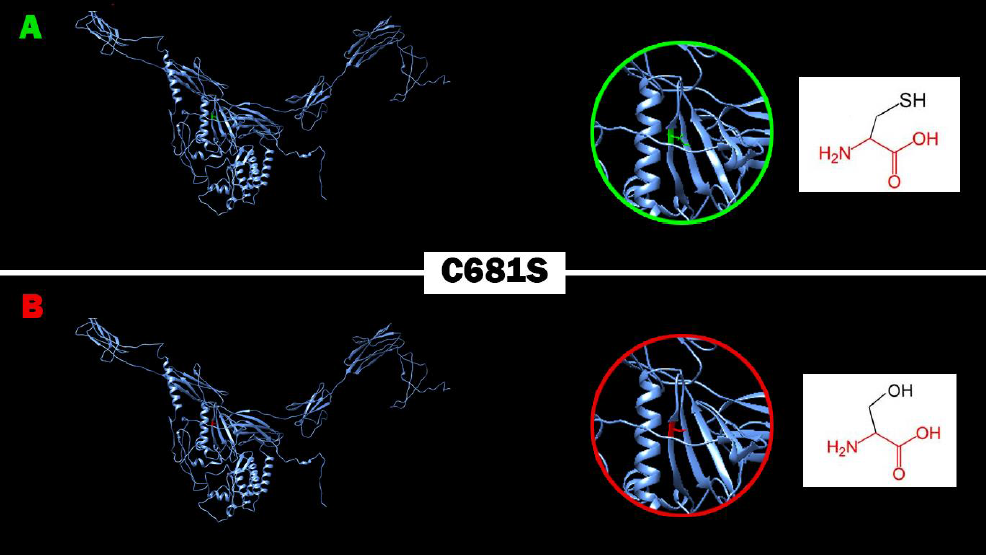
Effect of C681 (rs887807138) SNP on protein structure in which Cysteine (Green residue) mutated into Serine (Red residue) at position 681.

**Figure (4):**
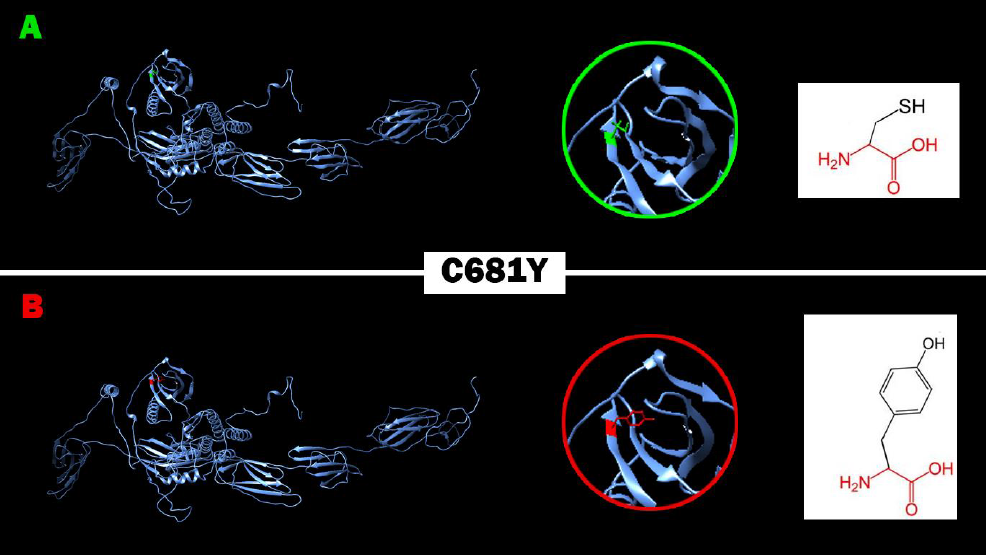
Effect of C681Y (rs887807138) SNP on protein structure in which Cysteine (Green residue) mutated into Tyrosine (Red residue) at position 681.

**Figure (5):**
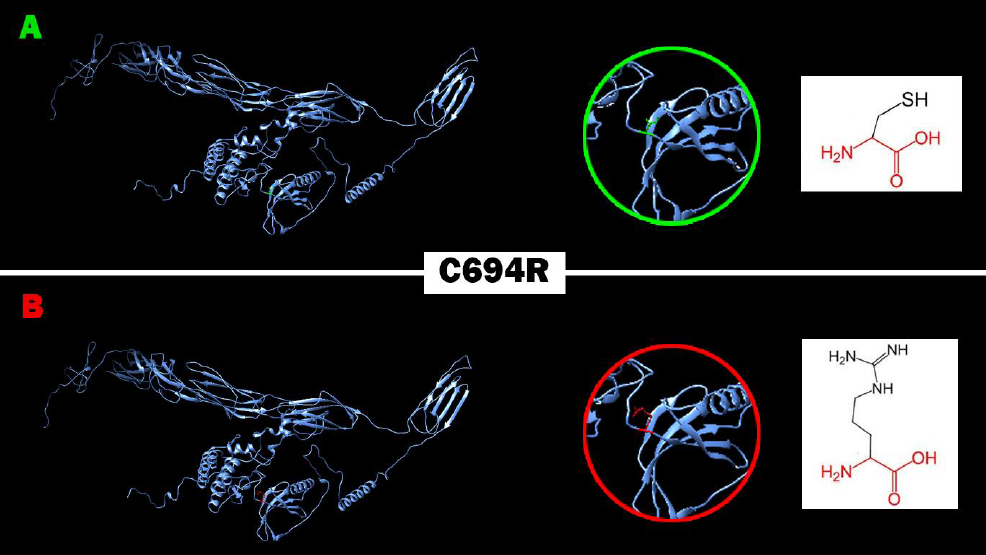
Effect of C694R (rs1221467960) SNP on protein structure in which Cysteine (Green residue) mutated into Arginine (Red residue) at position 694.

**Figure (6):**
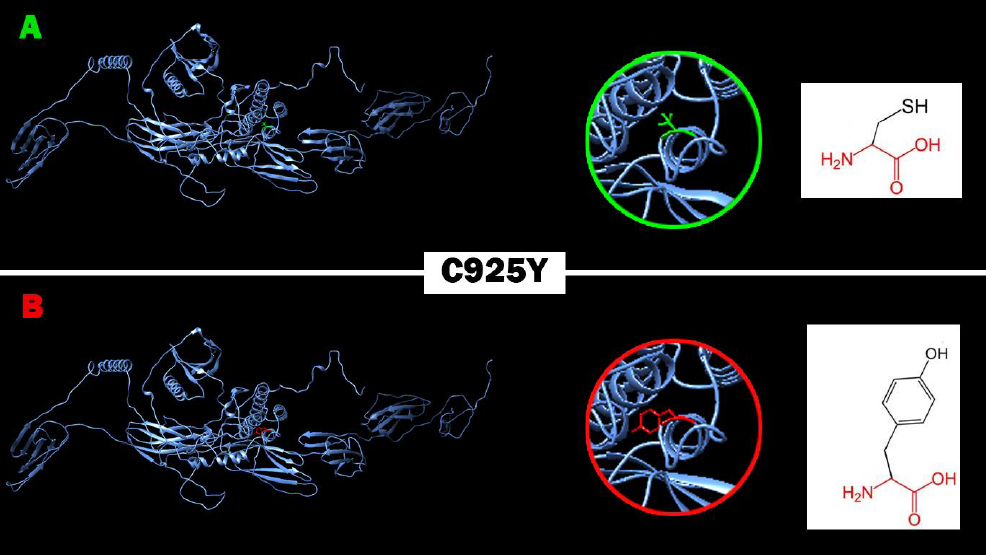
Effect of C925Y (rs1361689773) SNP on protein structure in which Cysteine (Green residue) mutated into Tyrosine (Red residue) at position 925.

**Figure (7):**
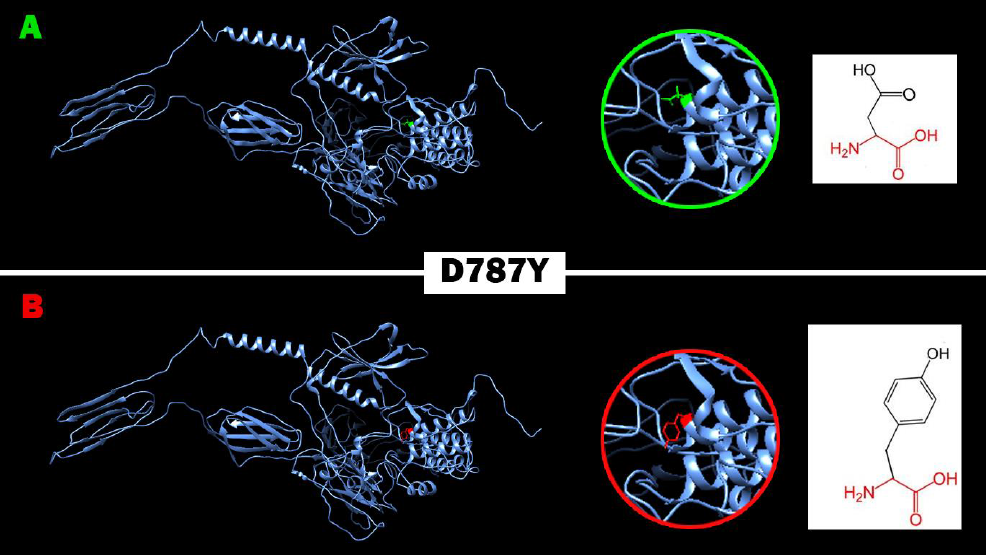
Effect of D787Y (rs1474119276) SNP on protein structure in which Aspartic acid (Green residue) mutated into Tyrosine (Red residue) at position 787.

**Figure (8):**
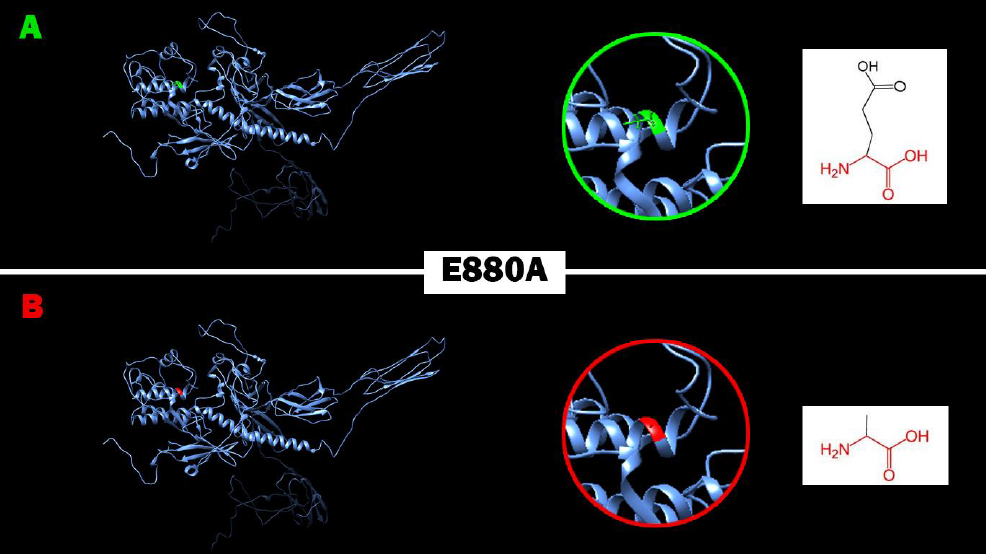
Effect of E880A (rs776952835) SNP on protein structure in which Glutamic acid (Green residue) mutated into Alanine (Red residue) at position 880.

**Figure (9):**
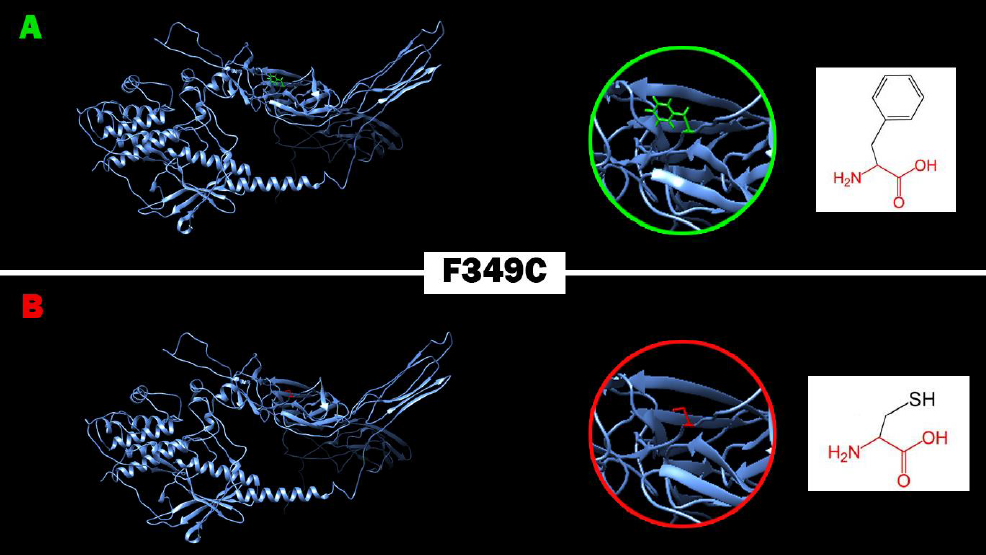
Effect of F349C (rs537215681) SNP on protein structure in which Phenylalanine (Green residue) mutated into Cysteine (Red residue) at position 349.

**Figure (10):**
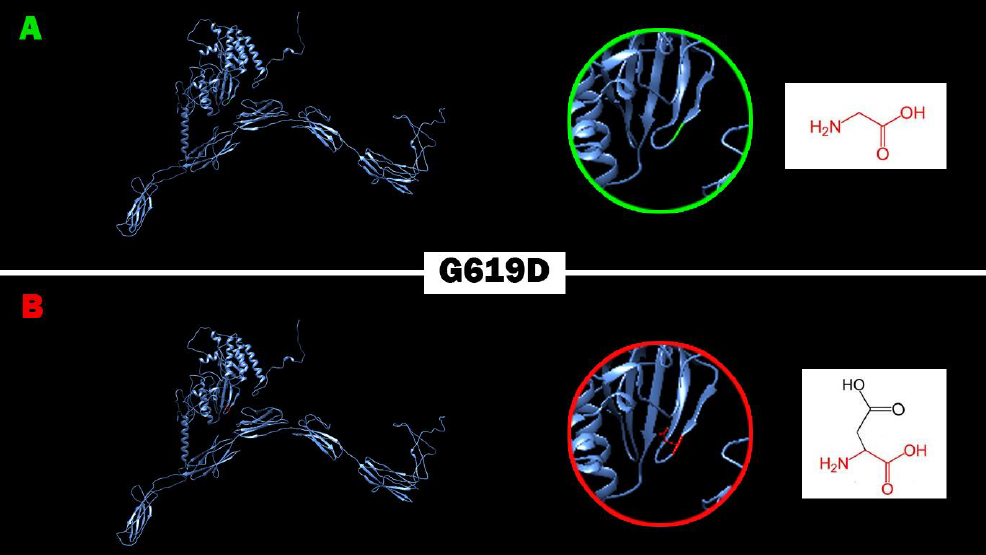
Effect of G619D (rs1442266284) SNP on protein structure in which Phenylalanine (Green residue) mutated into Cysteine (Red residue) at position 619.

**Figure (11):**
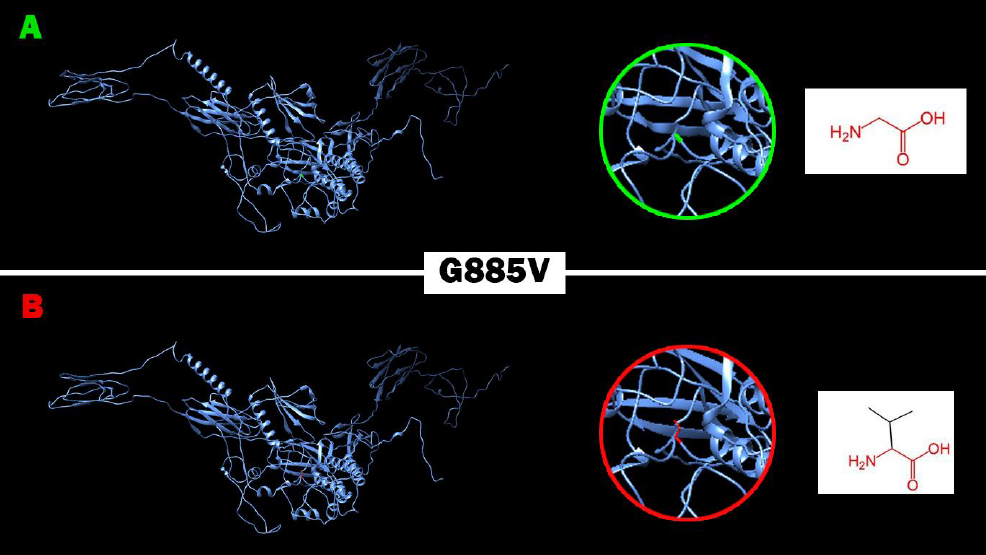
Effect of G885V (rs1442266284) SNP on protein structure in which Glycine (Green residue) mutated into Valine (Red residue) at position 885.

**Figure (12):**
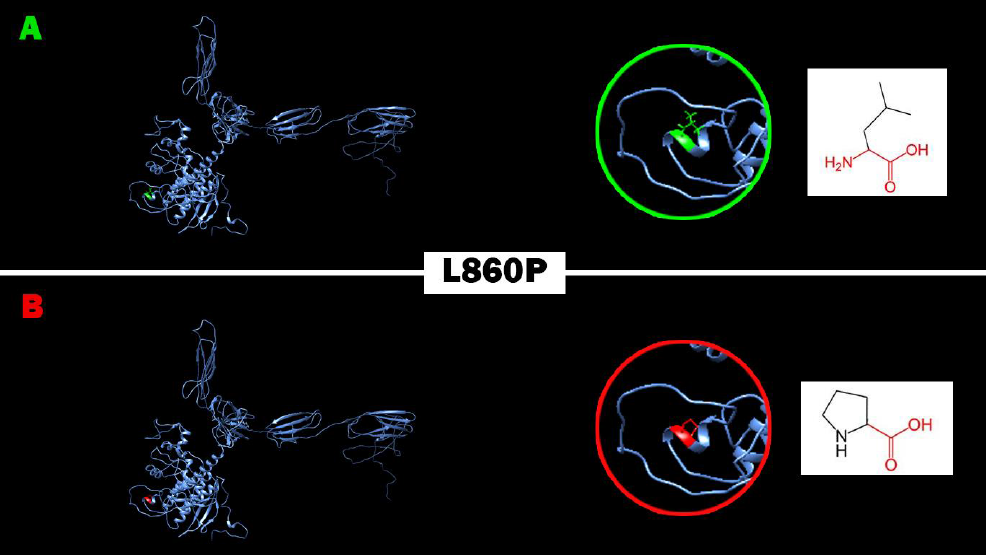
Effect of L860P (rs774360726) SNP on protein structure in which Leucine (Green residue) mutated into Proline (Red residue) at position 860.

**Figure (13):**
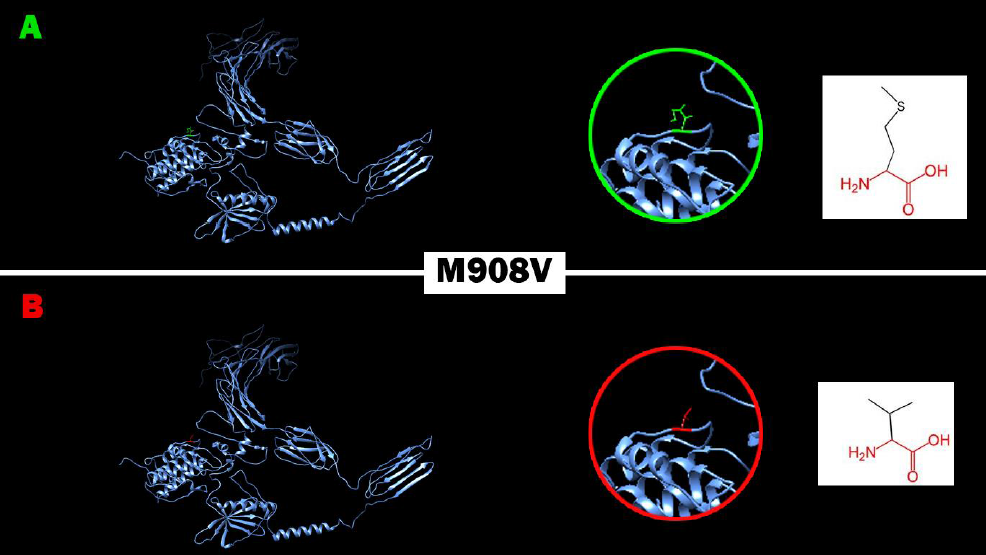
Effect of M908V (rs200909894) SNP on protein structure in which Methionine (Green residue) mutated into Valine (Red residue) at position 908.

**Figure (14):**
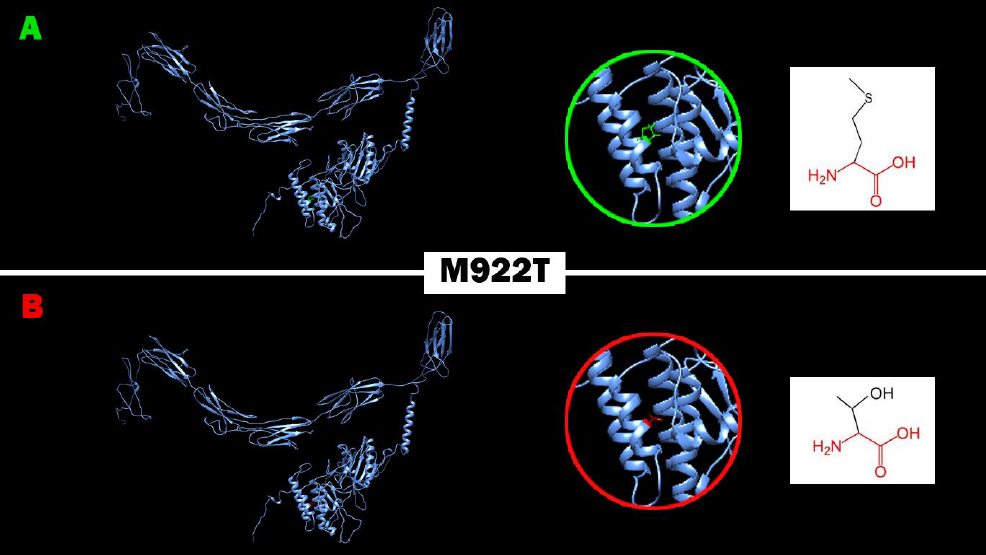
Effect of M922T (rs974980406) SNP on protein structure in which Methionine (Green residue) mutated into Threonine (Red residue) at position 922.

**Figure (15):**
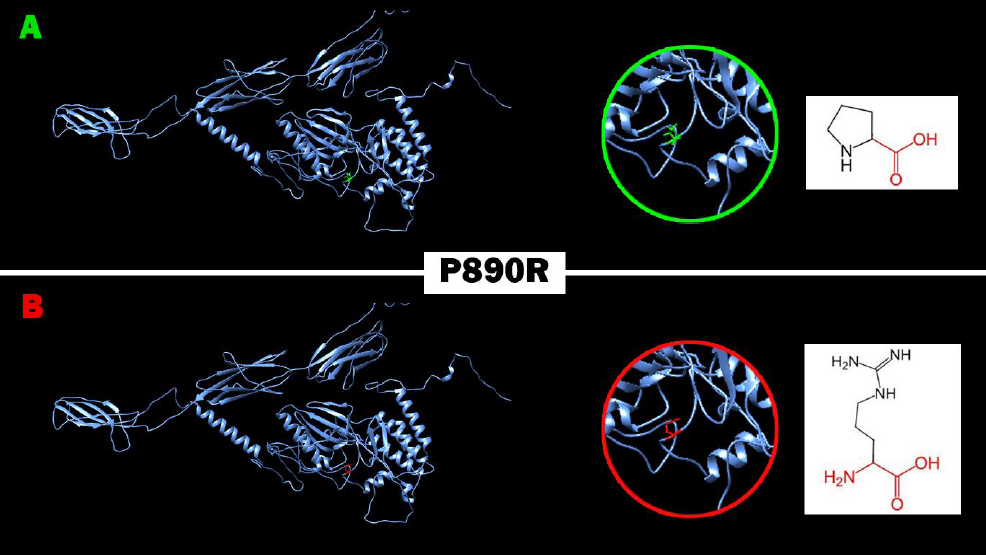
Effect of P890R (rs142796469) SNP on protein structure in which Proline (Green residue) mutated into Arginine (Red residue) at position 890.

**Figure (16):**
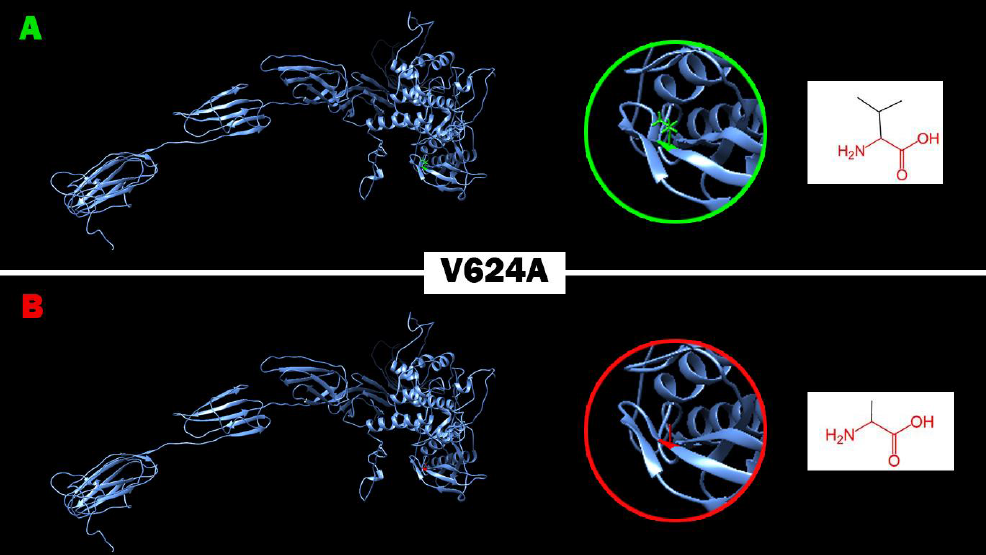
Effect of V624A (rs142796469) SNP on protein structure in which Valine (Green residue) mutated into Alanine (Red residue) at position 624.

**Figure (17):**
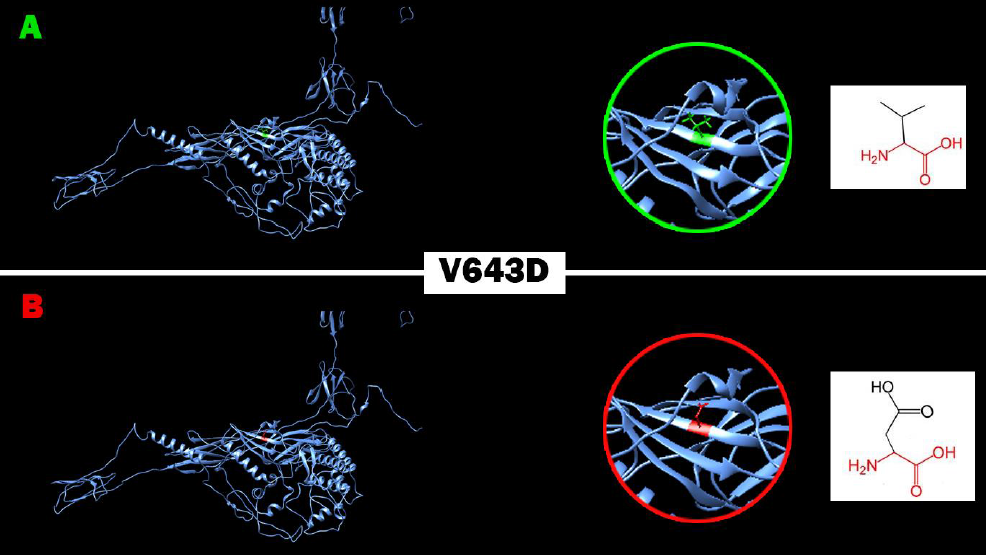
Effect of V643D (rs1276477642) SNP on protein structure in which Valine (Green residue) mutated into Aspartic acid (Red residue) at position 643.

**Figure (18):**
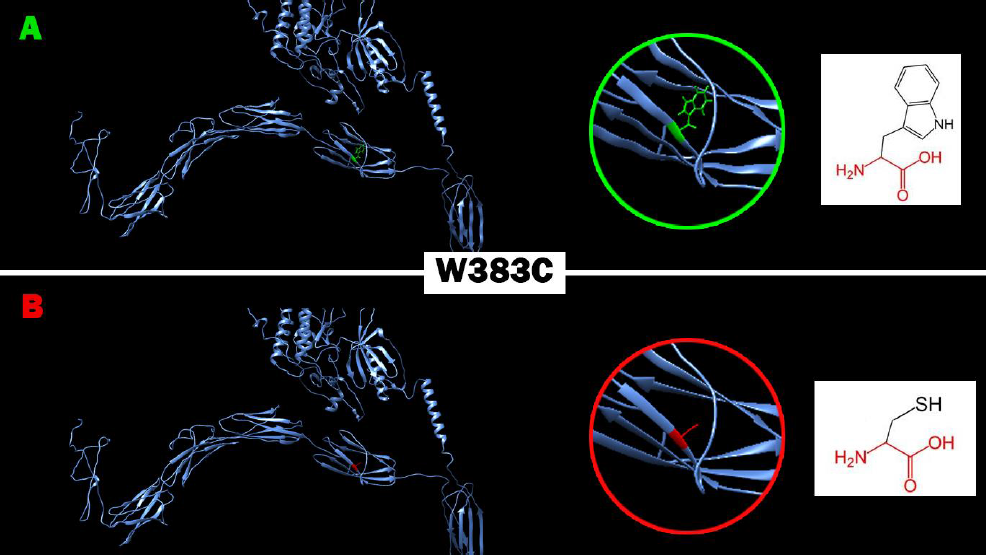
Effect of W383C (rs1041412646) SNP on protein structure in which Tryptophan (Green residue) mutated into Cysteine (Red residue) at position 383.

**Figure (19):**
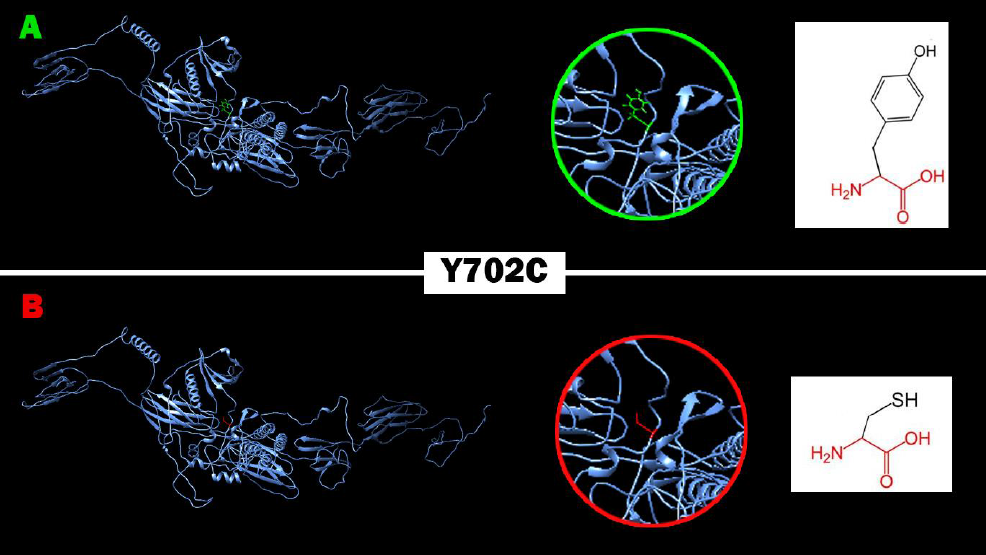
Effect of Y702C (rs774511256) SNP on protein structure in which Tyrosine (Green residue) mutated into Cysteine (Red residue) at position 702.

**Figure (20):**
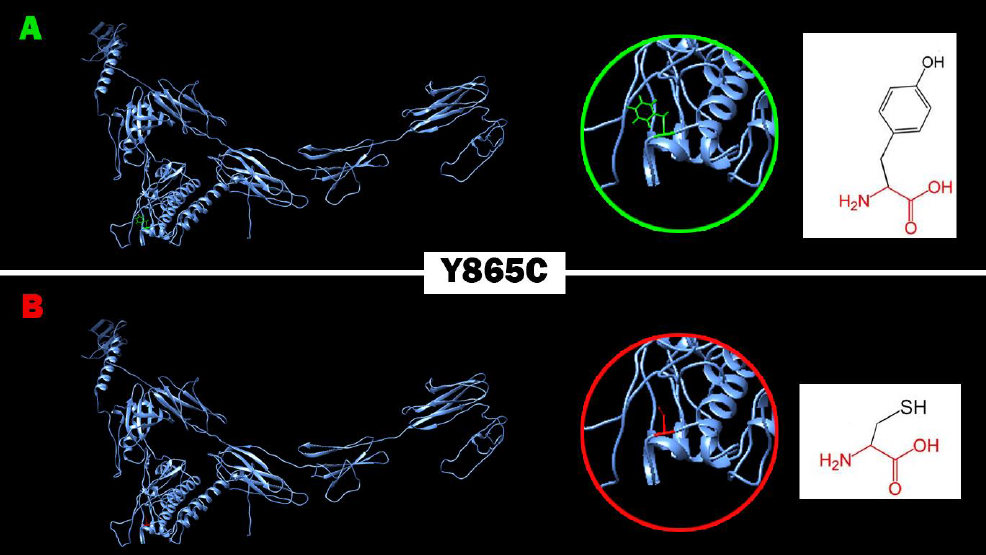
Effect of Y865C (rs759638195) SNP on protein structure in which Tyrosine (Green residue) mutated into Cysteine (Red residue) at position 865.

**Figure (21):**
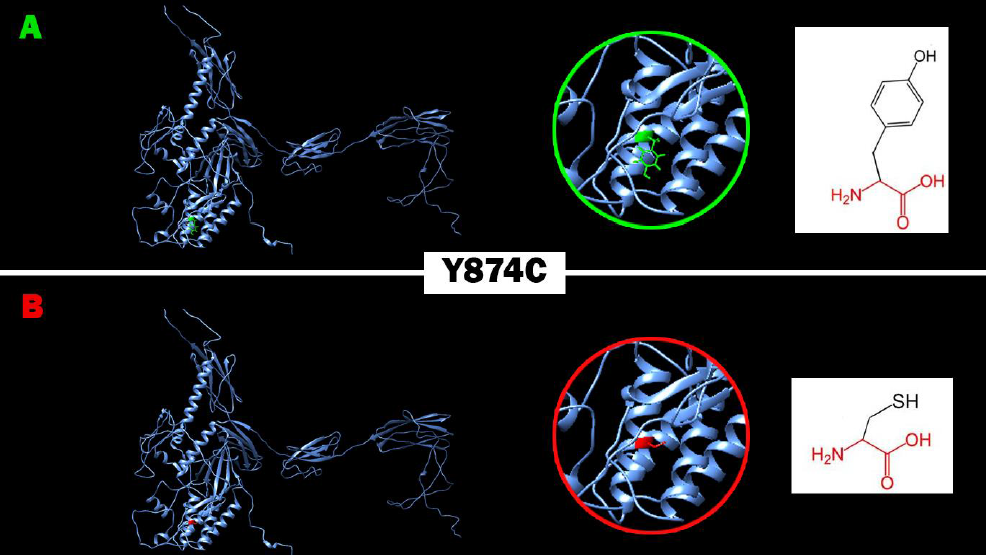
Effect of Y874C (rs868802040) SNP on protein structure in which Tyrosine (Green residue) mutated into Cysteine (Red residue) at position 874.

### SNPs effect on 3’UTR Region in *FLT3* gene using PolymiRTS Database

**Table (5):**
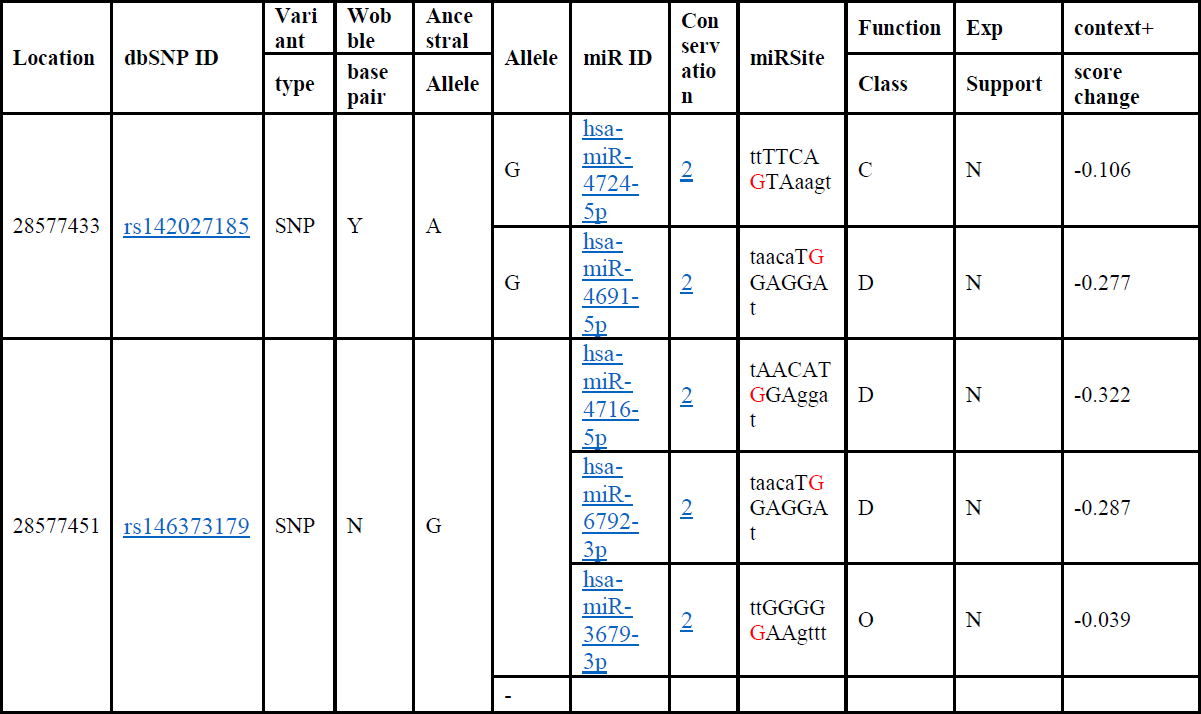
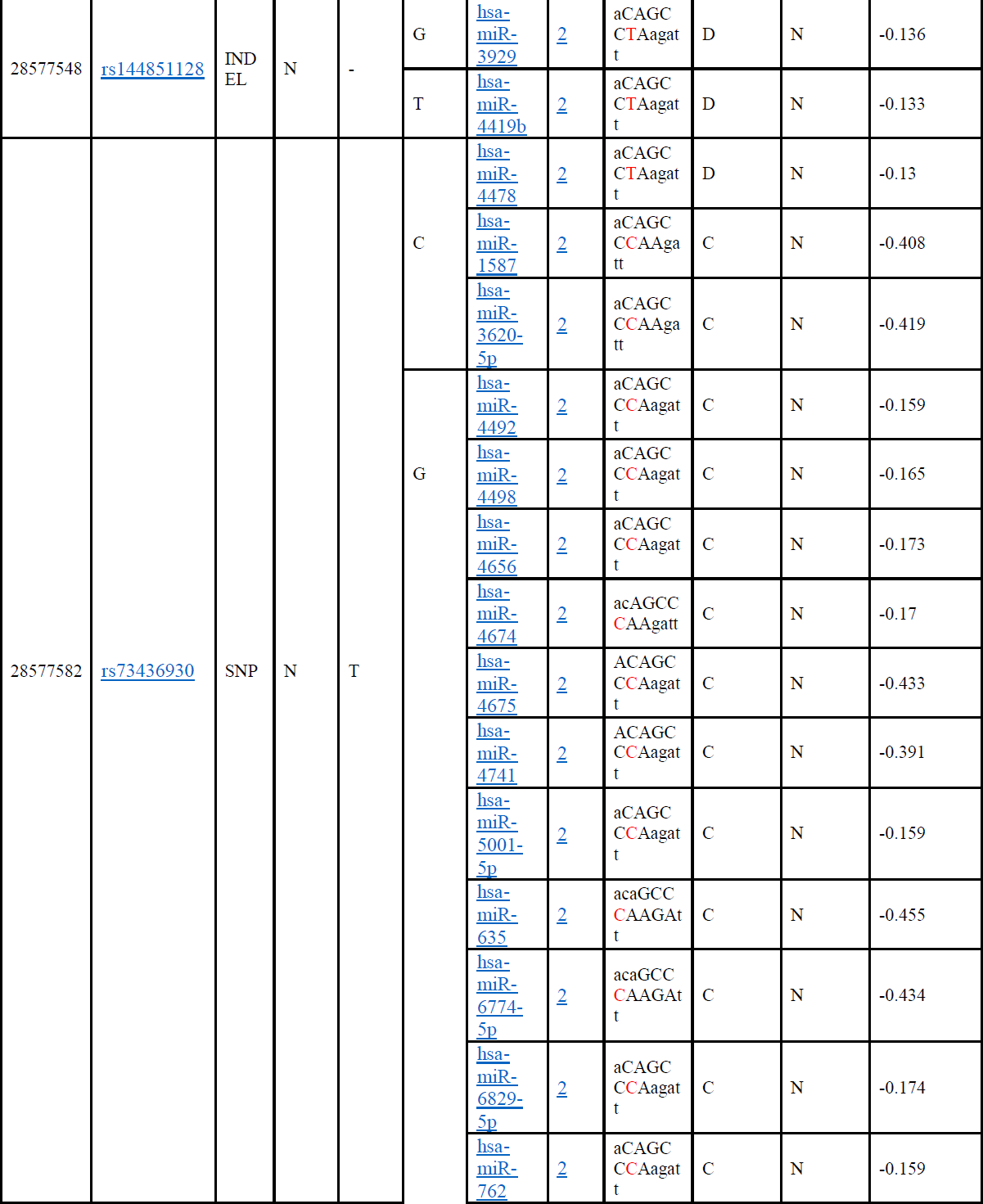
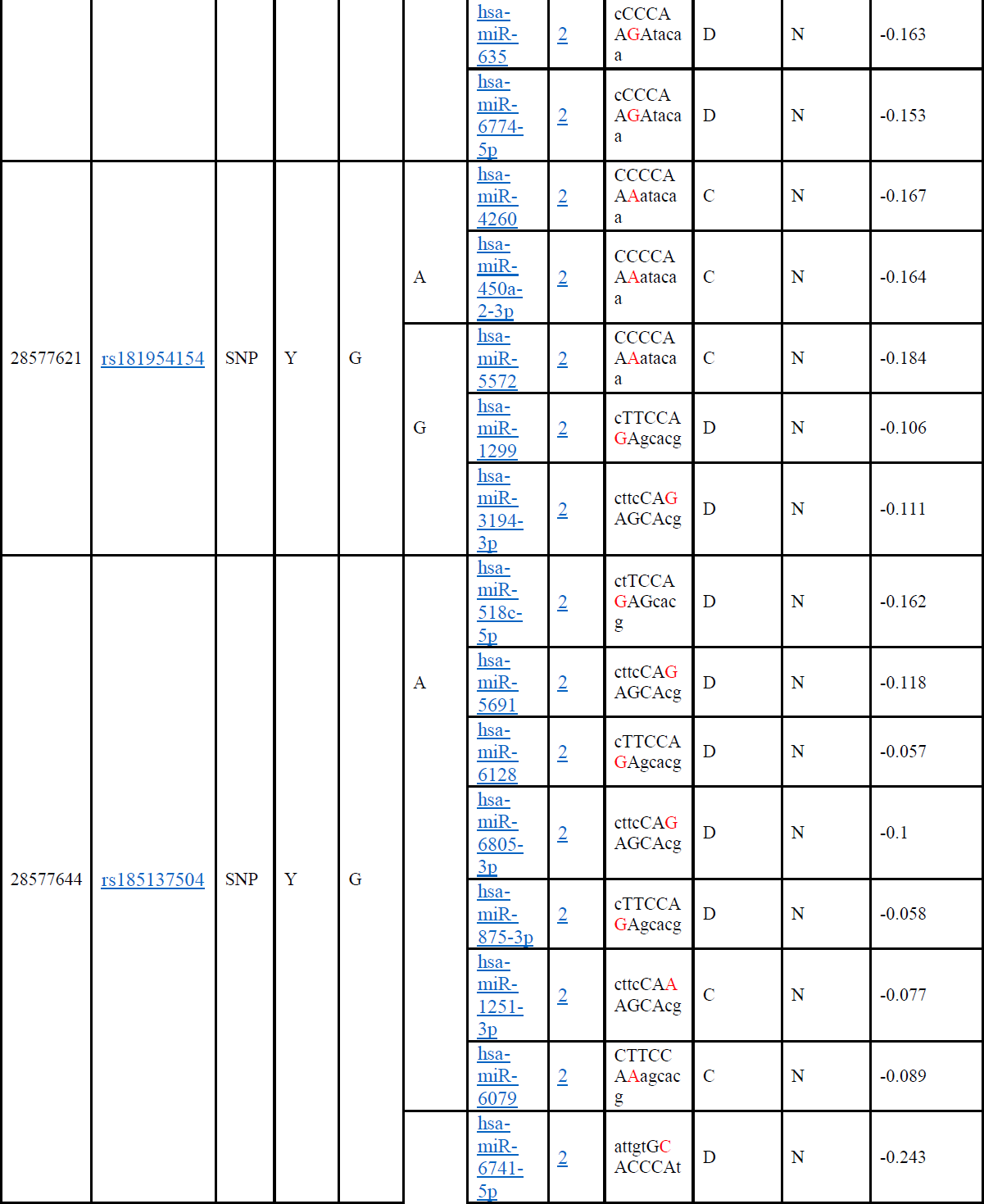
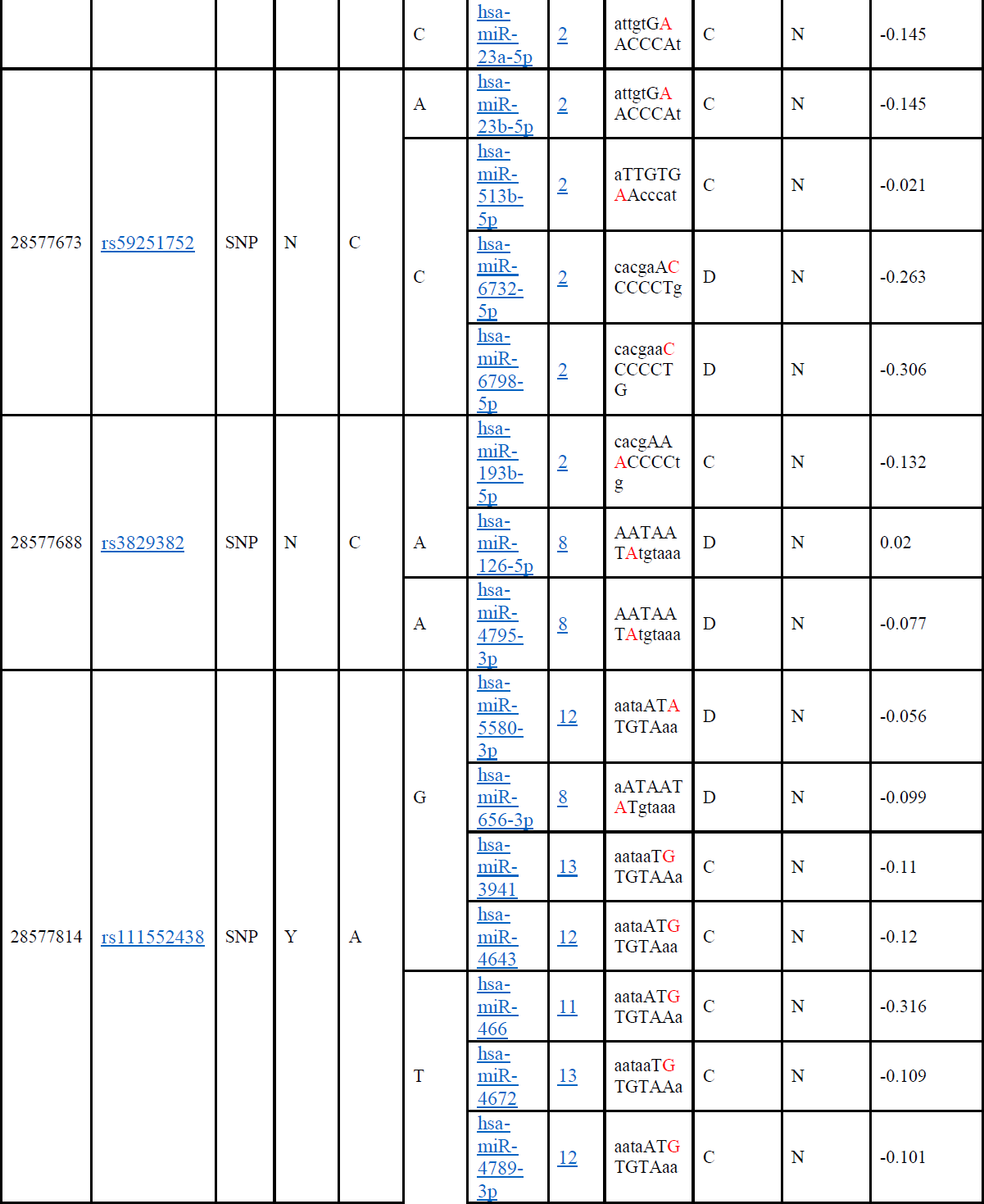
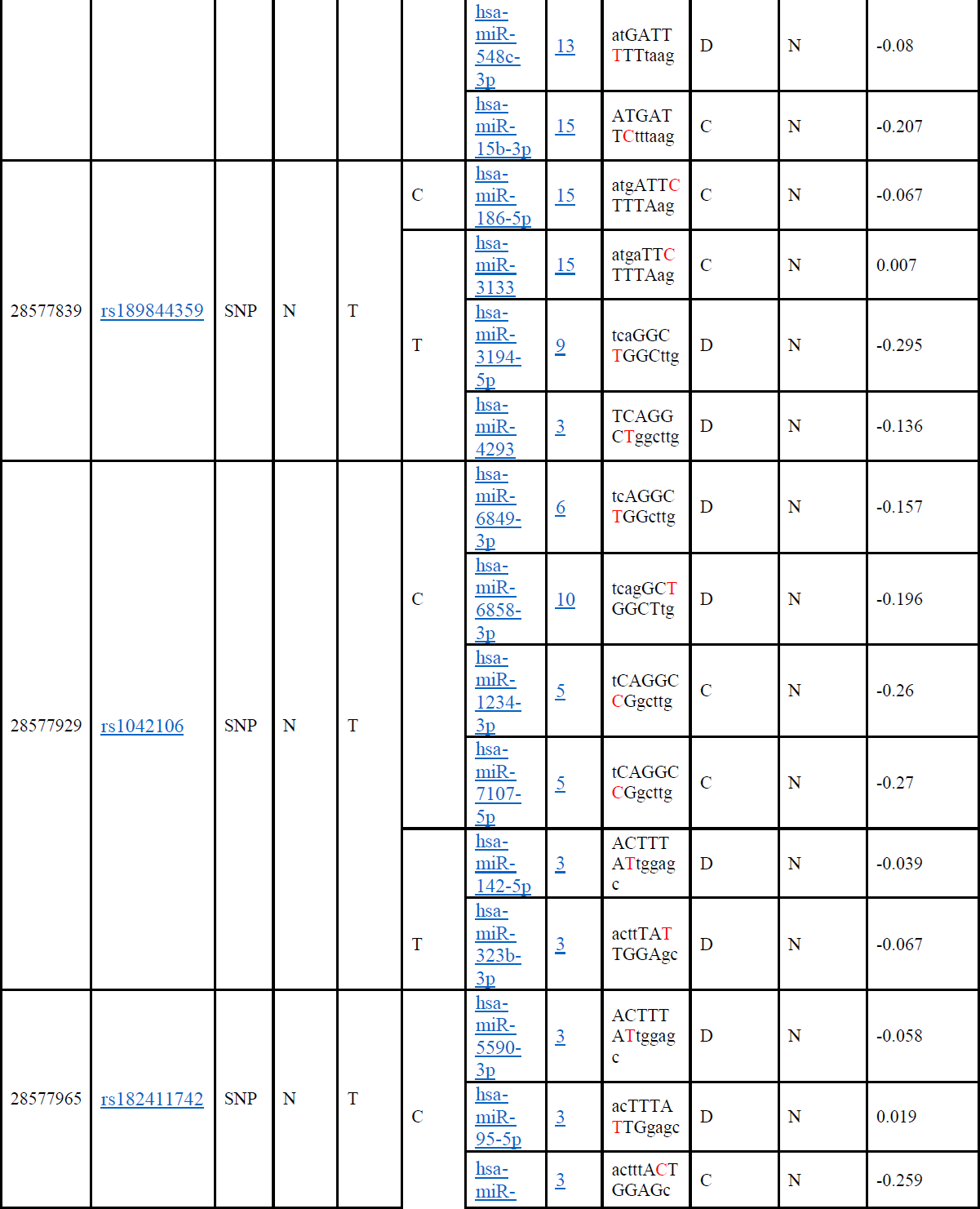
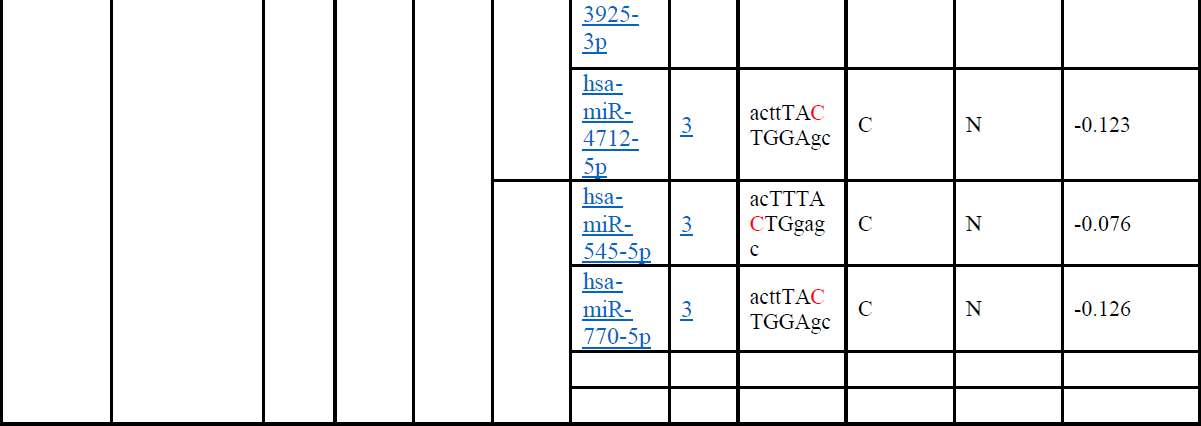
prediction of SNPs sites in NPM1 gene at the 3’UTR Region (miRNA binding sites) using PolymiRTS DB

### *FLT3* predicted gene functions and interactions with its appearance in network and genome using GeneMANIA software

**Table (3):** The *FLT3* gene functions and its appearance in network and genome:

***FLT3* predicted gene functions and interactions with its appearance in network and genome using GeneMANIA software:** **Table (3):** The *FLT3* gene functions and its appearance in network and genome:
| Function | FDR | Genes in network | Genes in genome |
| --- | --- | --- | --- |
| peptidyl-tyrosine modification | 3.45924E-05 | 7 | 252 |
| receptor signaling pathway via STAT | 3.45924E-05 | 6 | 138 |
| peptidyl-tyrosine phosphorylation | 3.45924E-05 | 7 | 227 |
| regulation of peptidyl-tyrosine phosphorylation | 0.000187402 | 6 | 192 |
| protein tyrosine kinase activity | 0.000212823 | 5 | 102 |
| extrinsic component of membrane | 0.000212823 | 6 | 210 |
| phosphatidylinositol 3-kinase regulator activity | 0.001325395 | 3 | 15 |
| receptor signaling pathway via JAK-STAT | 0.015560763 | 4 | 116 |
| lipid phosphorylation | 0.018813919 | 3 | 38 |
| phosphatidylinositol metabolic process | 0.044344056 | 4 | 160 |
| protein tyrosine kinase binding | 0.049960657 | 3 | 56 |

**Table (4):**
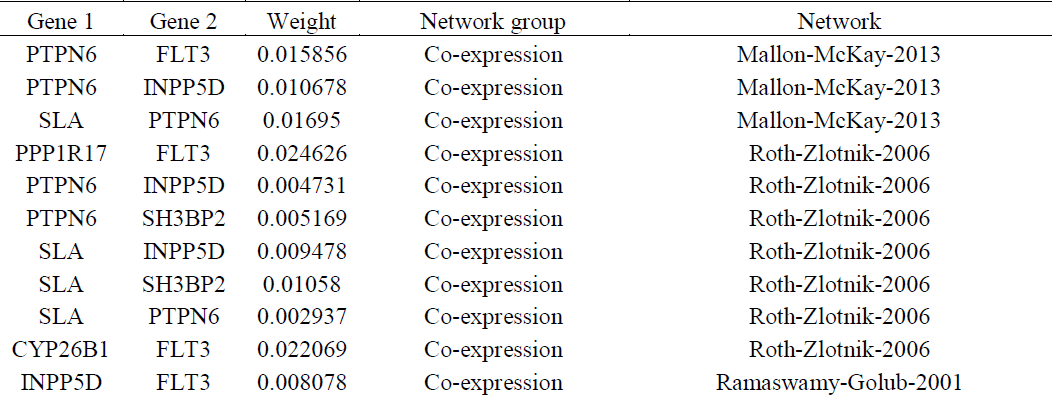
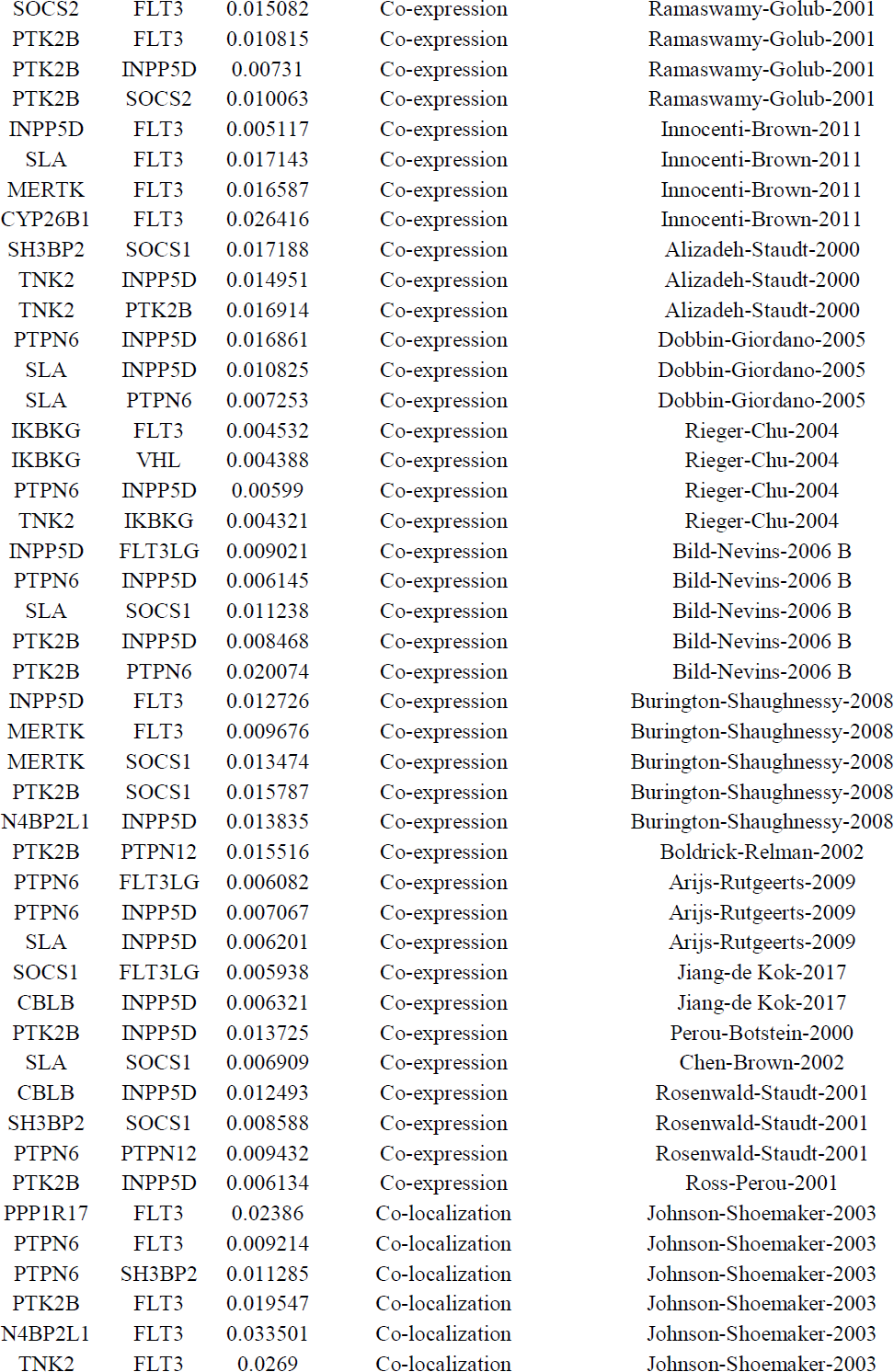
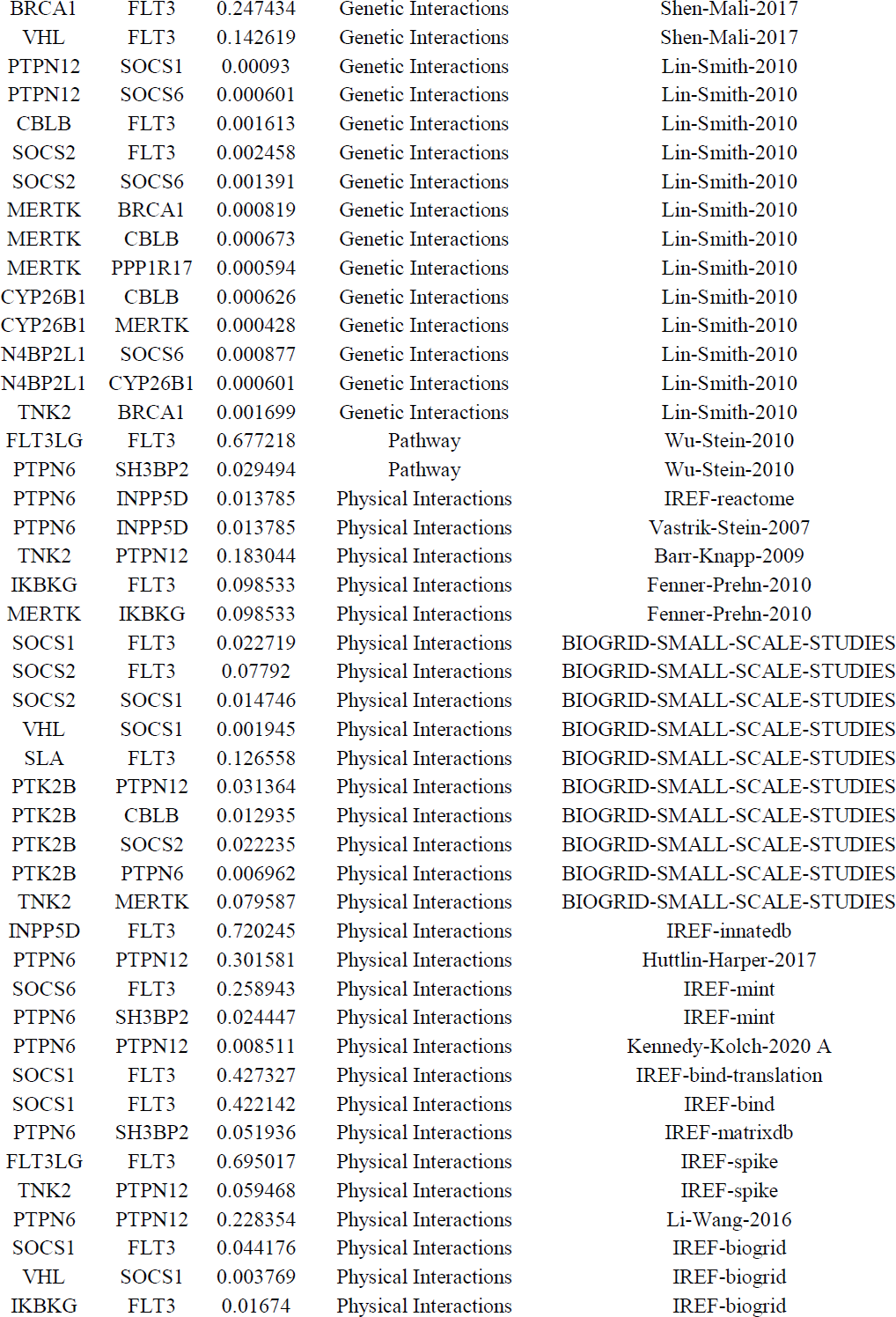
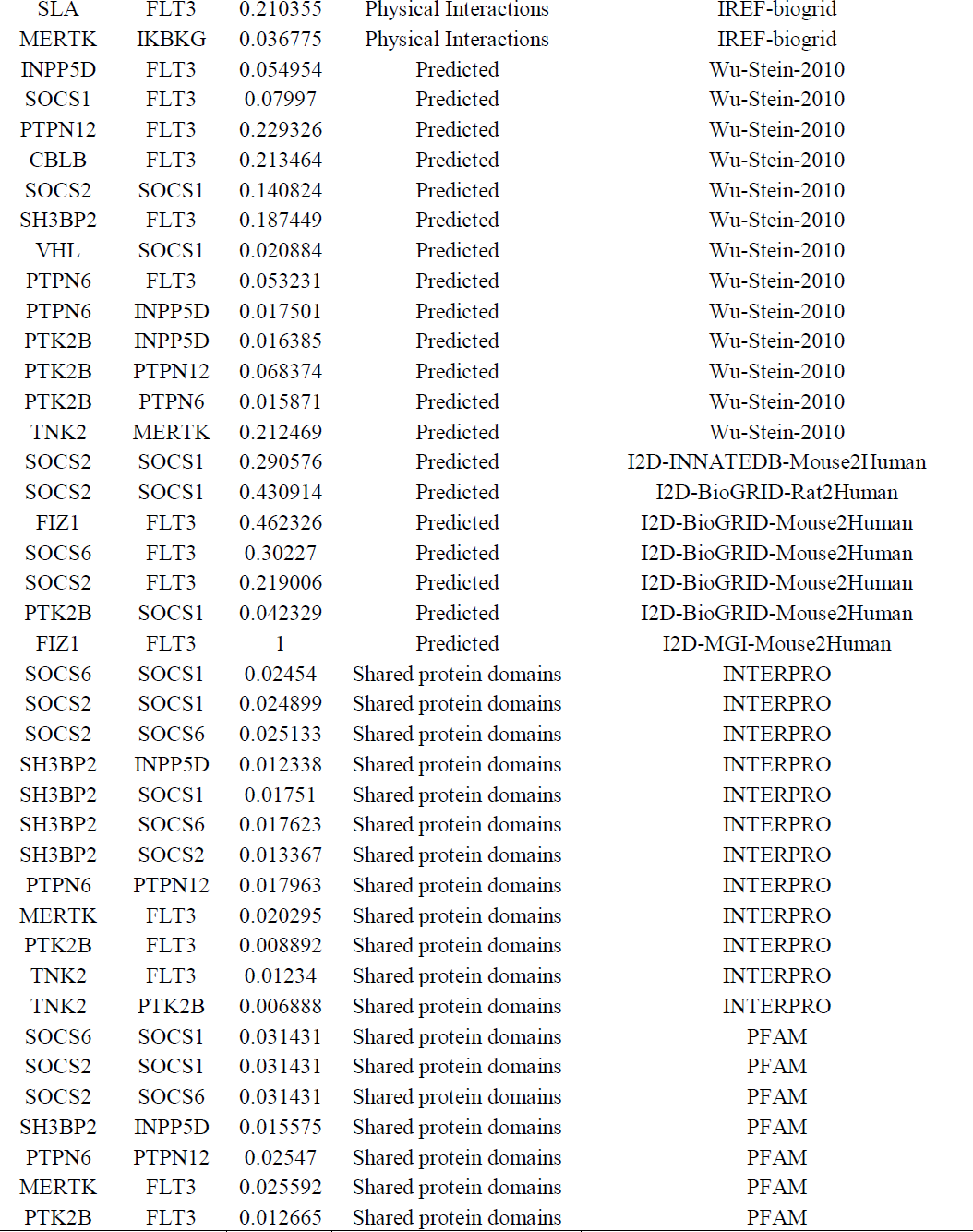
the gene co-expressed, share protein domain and Physical Interactions with *FLT3* gene network: Interactions of *FLT3* gene with other Functional Genes predicted by GeneMania:

**Figure (8):**
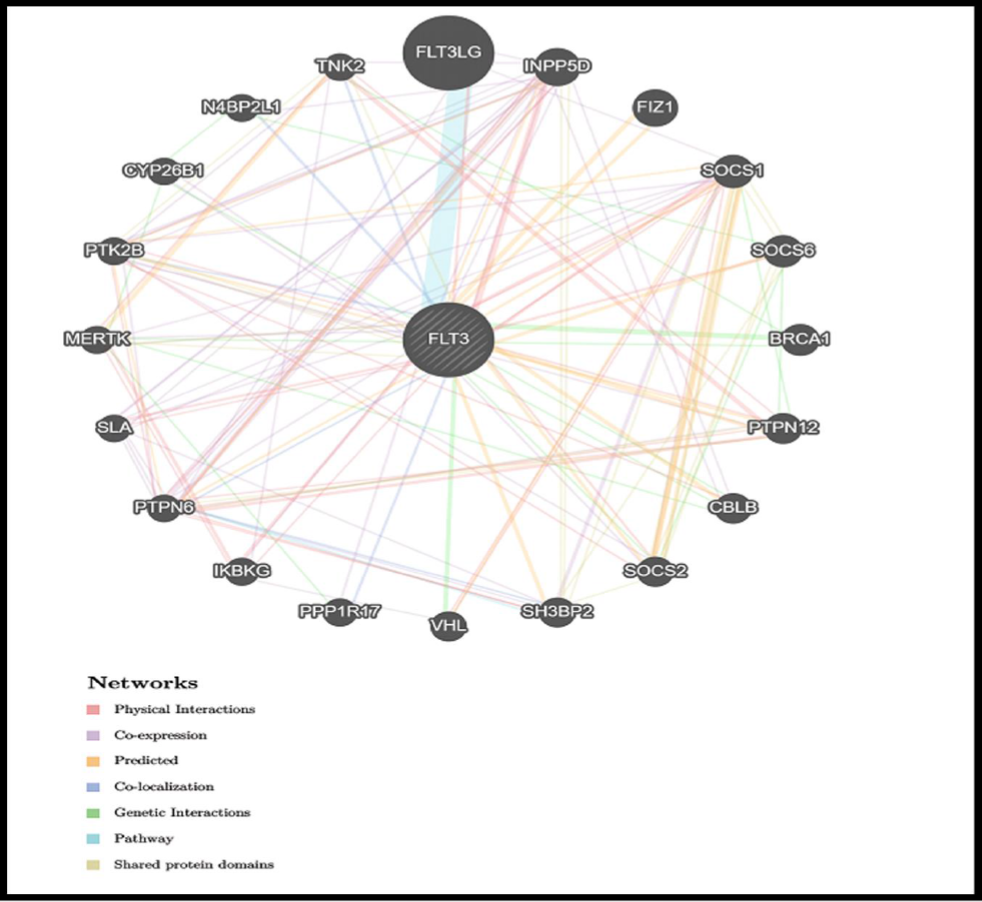
*FLT3* Gene Interactions and network predicted by GeneMania.

## DISCUSSION

In this study we revealed 20 novel SNPs out of 21 damaging SNPs in *FLT3* gene. Most damaging missense mutations (in coding region of gene) are detected by using different softwares.

E858K mutation is only SNP found in previous studies in AML patient as acquired mutation and in relapsed patients (25-27). The novel SNPs will be discussed by computational analysis study (C231Y, F349C, W383C, G619D, V624A, V643D, C681Y, C681S, C694R, Y702C, D787Y, L860P, Y865C, Y874C, E880A, G885V, P890R, M908V, M922T, C925Y).

C231Y, C681Y and C925Y mutations of a Cysteine into a Tyrosine at position 231, 681 and 925 respectively. The original wild-type residue and newly introduced mutant residue often differ in these properties. Regarding residue size the wild-type is smaller than mutant residue and concerning hydrophobicity the wild-type residue is more hydrophobic than the mutant residue which could cause loss of hydrophobic interaction in the core of the protein. Moreover, the wild-type residue was buried in the core of the protein at conserved positions. Based on this conservation information these mutations are probably damaging to the protein.

Y702C, Y865C and Y874C mutations of a Tyrosine into a Cysteine at position 702, 865 and 874 respectively affecting amino acid properties. The mutant residue is smaller and more hydrophobic than the wild-type residue. The mutation will cause loss of hydrogen bonds in the core of the protein and as a result disturb correct folding. Moreover, this will cause a possible loss of external interactions.

D787Y, V643D, C694R, G619D and G885V changing amino acids properties. Mutations affect hydrophobicity in which mutant residue is more hydrophobic than the wild-type residue; they affect size too as mutant residue is bigger than the wild-type residue. The wild-type residue of G885V was buried in the core of the protein probably will not fit. The torsion angles for this residue are unusual, only glycine is flexible enough to make these torsion angles, mutation into another residue will force the local backbone into an incorrect conformation and will disturb the local structure. On other hand, D787Y affecting charge in which wild type with NEGATIVE charge become NEUTRAL in opposite to V643D and G619D which changed from NEUTRAL to NEGATIVE. On other hand, C694R wild-type residue charge was NEUTRAL changed into mutant residue with POSITIVE charge.

L860P, W383C and V624A mutations located in conserved residues. The mutant residues are smaller than the wild-type residue. The former one will cause an empty space in the core of the protein and may cause a possible loss of external interactions. Moreover, the mutated residue affects the main activity of the protein and disturb its function.

Both M908V and M922T mutations affecting residue size in which mutant residue become smaller than the wild-type residue. Moreover, these mutations will cause an empty space in the core of the protein. Beside that M922T mutation will cause loss of hydrophobic interactions in the core of the protein.

E880A mutation of a Glutamic Acid into Alanine at position 880. There is a difference in charge, size and hydrophobicity between the wild-type and mutant amino acid. Regarding charge, the charge of the buried wild-type residue is lost by this mutation it turns from wild NEGATIVE residue into NEUTRAL mutant residue. On other hand, the size changed to be smaller mutant residue than the wild-type residue which may cause an empty space in the core of the protein. Moreover, the hydrophobicity of the wild-type and mutant residue differs. The mutant residue is more hydrophobic than the wild-type residue which may cause loss of hydrogen bonds in the core of the protein and as a result disturb correct folding.

P890R mutation located in very conserved residue is usually damaging for the protein. Based on this conservation information this mutation is probably damaging to the protein. The mutant residue is bigger than the wild-type residue which have NEUTRAL charge turning into POSITIVE mutant residue. The wild-type residue is more hydrophobic than the mutant residue. On other hand, C681S mutation also located in very conserved residue but may cause loss of hydrophobic interactions in the core of the protein.

Unlike Other mutations, the wild-type residue of F349C mutation is predicted to be located in its preferred secondary structure, a β-strand. Therefore, the local conformation will be slightly destabilized. On other hand, wild-type residue located in highly conserved position so mutation affects the activity of the protein and may disturb its function. Moreover, this mutation affect amino acid size as the mutant residue is smaller which might lead to loss of interactions.

In 3’UTR, out of the 148 SNPs there were 12 SNPs found to have an effect on it. 69 functional classes were predicted among the 12 SNPs; 31 alleles disrupted a conserved miRNA site and 37 derived alleles creating a new site of miRNA and this might result in deregulation of the gene function.

Genmania predicts that *FLT3* has peptidyl-tyrosine modification, peptidyl-tyrosine phosphorylation, protein tyrosine kinase activity, receptor signaling pathway via JAK-STAT, receptor signaling pathway via STAT, regulation of peptidyl-tyrosine phosphorylation activities. As bioinformatics analysis results possibly have a valuable outcome for patient suffering from Acute Myeloid Leukemia (AML) with a mutation in *FLT3* protein but in-vivo/in-vitro analysis is highly suggested. These findings could be valuable in diagnosis and treatment of AML patients.

## CONCLUSION

This study revealed 20 damaging SNPs considered to be novel nsSNP in *FLT3* gene that leads to AML, by using different algorithms. Additionally, 69 functional classes were predicted in 12 SNPs in the 3’UTR, among them, 31 alleles disrupted a conserved miRNA site and 37 derived alleles created a new site of miRNA. This might result in the de regulation of the gene function. These results could be valuable for molecular studying, diagnosis and treatment of AML patients.

## ACKNOWLEDGMENT

The authors wish to acknowledgment the enthusiastic cooperation of Alneelain Stem Cell Center & Africa City of Technology, Khartoum - Sudan.

## DATA AVAILABILITY

All relevant data used to support the findings of this study are included within the manuscript and supplementary information files.

